# Raman spectroscopy-based measurements of single-cell phenotypic diversity in microbial communities

**DOI:** 10.1101/2020.05.21.109934

**Authors:** Cristina García-Timermans, Ruben Props, Boris Zacchetti, Myrsini Sakarika, Frank Delvigne, Nico Boon

## Abstract

Microbial cells experience physiological changes due to environmental change, such as pH and temperature, the release of bactericidal agents, or nutrient limitation. This, has been shown to affect community assembly and other processes such as stress tolerance, virulence or cell physiology. Metabolic stress is one such physiological changes and is typically quantified by measuring community phenotypic properties such as biomass growth, reactive oxygen species or cell permeability. However, community measurements do not take into account single-cell phenotypic diversity, important for a better understanding and management of microbial populations. Raman spectroscopy is a non-destructive alternative that provides detailed information on the biochemical make-up of each individual cell.

Here, we introduce a method for describing single-cell phenotypic diversity using the Hill diversity framework of Raman spectra. Using the biomolecular profile of individual cells, we obtained a metric to compare cellular states and used it to study stress-induced changes. First, in two *Escherichia coli* populations either treated with ethanol or non-treated. Then, in two *Saccharomyces cerevisiae* subpopulations with either high or low expression of a stress reporter. In both cases, we were able to quantify single-cell phenotypic diversity and to discriminate metabolically stressed cells using a clustering algorithm. We also described how the lipid, protein and nucleic acid composition changed after the exposure to the stressor using information from the Raman spectra. Our results show that Raman spectroscopy delivers the necessary resolution to quantify phenotypic diversity within individual cells and that this information can be used to study stress-driven metabolic diversity in microbial communities.

**Importance:** Microbes that live in the same community respond differently to stress. This phenomemon is known as phenotypic diversity. Describing this plethora of expressions can help to better understand and manage microbial processes. However, most tools to study phenotypic diversity only average the behaviour of the community. In this work, we present a way to quantify the phenotypic diversity of single cells using Raman spectroscopy - a tool that can describe the molecular profile of microbes. We demonstrate how this tool can be used to quantify the phenotypic diversity that arises after the exposure of microbes to stress. We also show its potential as an ‘alarm’ system to detect when communities are changing into a ‘stressed’ type.

## Introduction

Monoclonal microbial populations can exhibit heterogeneous genetic expression, which underlies phenotypic differences between cells. Phenotypic diversity has been shown to increase population survival or fitness in a changing environment and allows microorganisms to divide tasks and organize as a group. This differential gene expression can arise due to environmental pressure, stochastic events, periodic oscillations or cell-to-cell interactions (Ackermann, 2015; Altschuler & Wu, 2010; Avery, 2006). When a deviation from optimal growth conditions occurs such as changes in temperature, pH, nutrients salts and/or oxygen levels, a stress response is triggered in microorganisms (both prokaryotes and eukaryotes), resulting in a biochemical cascade to promote stress tolerance, virulence or other physiological changes. These strategies can result in enhanced survival, virulence, cross-protection or cell death (Ron, 2013; Święciło, 2016; Wesche et al., 2009). Usually, microorganisms show mixed behavioural strategies, maximizing the chances of survival (Lowery et al., 2017), making phenotypic diversity a crucial characteristic of stress-driven phenotypes. However, cellular stress is often measured at the community level using bulk technologies, such as cell concentration, quantity of reactive oxygen species (ROS), cell permeability or protein content. While these methods reveal important information, they provide the average information for the whole population, failing to describe cell-to-cell variability and bet-hedging strategies (Veening et al., 2008). To better understand stress-driven changes, single cell technologies must be used.

There are several single cell technologies available to study the response of individual cells to stress. For example, fluorescent labels that tag certain cellular functions (membrane potential, intracellular enzyme activity, a stress reporter) can be used in combination with flow cytometry (Delvigne et al., 2015; Porter et al., 1995) or imaging techniques (Benomar et al., 2015). Single-cell (multi)-omics opens the door to a very detailed understanding of the metabolism of individual cells, although it is a low-throughput technique that still presents many challenges in its accuracy (Bock et al., 2016). Raman spectroscopy is an alternative single-cell tool that can detect individual phenotypes without the use of fluorescent probes. It is an optical method in which the Raman scattering of a cell and/or particle is collected thereby generating a single-cell fingerprint that contains (semi)quantitative information on its constituent molecules, such as nucleic acids, proteins, lipids and carbohydrates. This technique has been used to study stress-induced phenotypic differences of the cyanobacterium *Synechocystis* sp. (Tanniche et al., 2020): the fingerprints of cells treated with different concentrations of acetate or NaCl and non-treated cells were differentiable using discriminant analysis of principal component analysis (PCA). Also, Teng and colleagues (Teng et al., 2016) found that *Escherichia coli* cells exposed to several antibiotics, alcohols and chemicals had distinct Raman fingerprints. However, there are currently no quantitative methods to describe phenotypic diversity in single cells using their Raman spectra.

A widely used set of metrics to quantify the diversity of microbial communities are Hill numbers, also known as the effective number of species, as they express in intuitive units the number of equally abundant species that are needed to give the same value of the diversity measure. Hill numbers respect other important ecological principles, such as the replication principle, that states that in a group with *N* equally diverse groups that have no species in common, the diversity of the pooled groups must be the *N* times the diversity of a single group (Chao et al., 2014; Daly et al., 2018). They are commonly used to quantify microbial diversity based on 16S rRNA sequencing techniques but have also been applied to flow cytometry yielding similar results (Props et al., 2016). However, phenotypic diversity at the single-cell level has not yet been described. This would require multiparametric information of individual cells, something Raman spectroscopy can provide.

Quantifying phenotypic diversity at the single-cell level could be useful to follow and manage stress in bioproduction: to maintain high bioproduction rates, it is important to find or create stress-tolerant organisms. For instance, in microbial production of alcohol (considered a sustainable alternative source for chemicals and fuels), one of the major limitations is the toxicity and/or growth inhibition caused by the alcohol that is produced. The alcohol increases the fluidity of the cell membrane and causes a disruption on the phospholipid components that inhibits growth and can lead to death. It also affects nutrient uptake and ion transport. Therefore, there have been efforts in evolutionary and synthetic engineering to increase alcohol tolerance in several organisms, for example, *E. coli* and *S. cerevisiae,* widely used in bioproduction (Jia et al., 2010).

We aim to quantify single-cell phenotypic diversity using Raman spectroscopy, based on the Hill diversity framework. We described the necessary steps to preprocess Raman spectra and demonstrate its integration into the Hill diversity framework. The necessary functionalities are also embedded in the open source MicroRaman package (https://github.com/CMET-UGent/MicroRaman). To illustrate the use of this method, we applied it in two popular strains in bioproduction. First, we compared an *E. coli* population in stress conditions (cultivated with ethanol) with a control population. Secondly, we separated two subpopulations of a *S. cerevisiae* culture that was under nutrient-limiting conditions using a GFP tag and analyzed them using Raman spectroscopy. In both cases, we show how the stress-induced single-cell phenotypic diversity can be quantified using the Raman spectra of the single cells, and how this information can be used to detect a shift in the phenotype of the population. Finally, we use this information to explain how the molecular profile of the cells changes after being exposed to the stressors.

## Materials and methods

### Data sets

The strains used and the incubation medium are described in Table 1. We did ~450 measurements in 4 axenic cultures using Raman spectroscopy. Samples were cultured at 28°C with 120 rpm orbital shaking. Each strain was re-cultivated via transferring 10% v/v of active culture in fresh liquid medium (described in table 1) every 24 to 48h for 2 months. Cultures were harvested by centrifugation at 6603 g for 5 min, washed with 0.1M phosphate buffer saline (PBS) and stored at −4°C until further use.

**Table 1:**
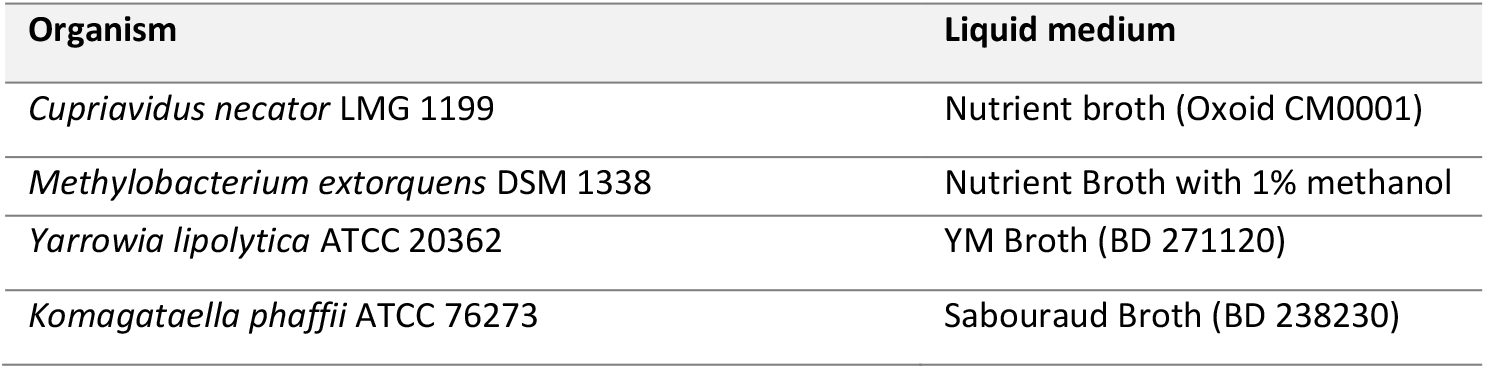
List of organisms and medium used to grow them

### Case studies: single-cell phenotypic diversity quantification in stress-induced phenotypes

To test the capacity of the single-cell phenotypic diversity (sc-D_2_) calculation to identify metabolic changes, we used two case studies. First, we studied two *E. coli* populations that had been grown together in different conditions: one was treated with ethanol while the other was not. Secondly, a *S. cerevisiae* culture was grown in nutrient limiting conditions, which resulted in differential expression of the chimeric stress reporter (tagged with eGFP). The two subpopulations (high expressing and low expressing eGFP) were isolated (Fig 1).

**Fig 1:**
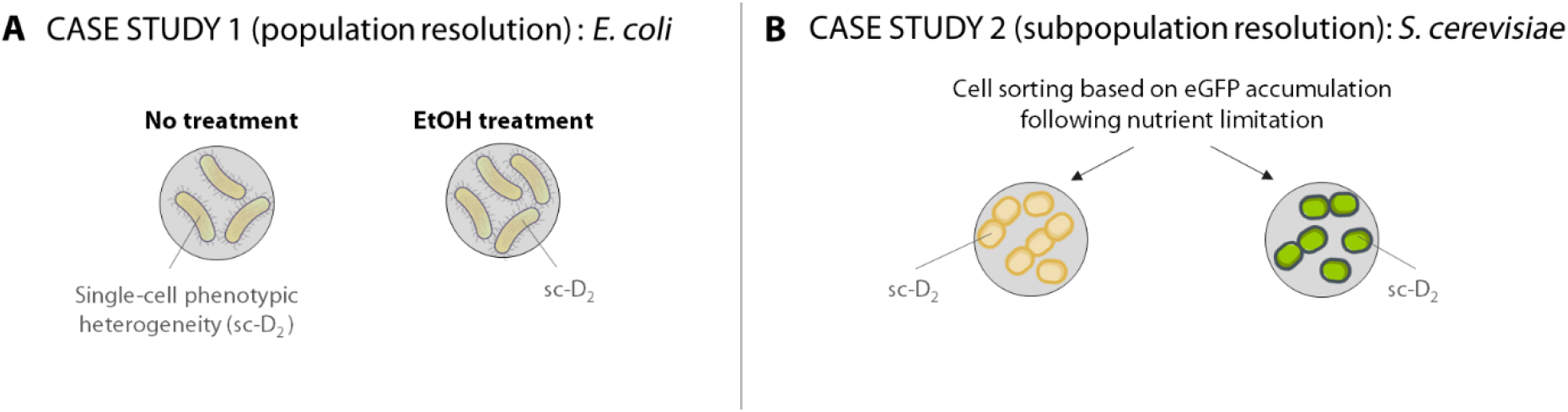
Overview of the case studies. A) Study of two *E. coli* populations grown separately with ethanol in the medium or non-treated. B) Two subpopulations were isolated from a *S. cerevisiae* culture based on the expression of the GFP marked chimeric stress reporter after nutrient limitation. The Raman spectra of single cells were used to calculate their phenotypic diversity (sc-D_2_).

#### Population resolution: *E. coli* exposed to ethanol

The dataset from Teng et al. 2016 was used to validate alpha and beta-diversity calculations. According to their manuscript, this dataset consists of Raman spectra of *Escherichia coli* in different time intervals (5, 10, 20, 30 and 60 min, 3 h and 5 h) after being cultured with different chemical stressors. We used the ethanol-treated samples and the controls to illustrate our point. The dataset consists of three biological replicates of the cell culture and measured 20 cells per replicate.

#### Subpopulation resolution: *S. cerevisiae* after nutrient limitation

The prototrophic haploid yeast strain *Saccharomyces cerevisiae* CENPK 113-7D was used in this study (Nijkamp et al., 2012). eGFP was produced under the control of a chimeric promoter composed of fragments of the *HSP26* and *GLC3* promoters. The promoter sequence was previously published (chimaera 2 in (Zid & O’Shea, 2014)). A synthetic construct containing the promoter, the eGFP gene and the G418 resistance marker was integrated in the genome via homologous recombination at the *uga1* site. The correct insertion was confirmed via PCR analysis and lack of growth on gamma-aminobutyrate (GABA) as the sole nitrogen source.

Samples were collected after 10 residence times in a continuous culture operated at D=0.1 h^-1^ in a 2-liter stirred-tank bioreactor with 1 liter operating volume. Defined yeast mineral medium containing 7.5 g l^-1^ was used (Verduyn et al., 1992). The culture temperature was maintained at 30° C, the stirrer speed at 1000 rpm and the air provision at 1 vvm. The culture pH was controlled at 5.0 through the automated addition of either 25% KOH or 25% M H_3_PO_4_.

Before cell sorting, samples were fixed in formaldehyde 4%, following the protocol from García-Timermans et al., 2018. Paraformaldehyde is known to preserve the Raman spectral features better than other fixatives, such as ethanol or glutaraldehyde (Read & Whiteley, 2015). Upon reaching steady-state in nutrient limited continuous culture, yeast population was sorted in two distinct subpopulations, i.e. the first one exhibited a high GFP content (high GFP) and the second one exhibiting a low GFP content (low GFP). Then, the high GFP and low GFP subpopulations were separated using Fluorescence-activated cell sorting (FACS). For this purpose, cell suspension collected from the bioreactor was diluted 10 times in PBS (ThermoFischer scientific, Belgium) and was further analyzed and sorted with a FACSaria (Becton Dickinson, Belgium). Cells have been collected following an enrichment sorting mode. Fractions containing 10^6^ cells of each subpopulation were collected. (Gating details used for cell sorting can be found in Supplementary Information).

### Raman spectroscopy

For the *S. cerevisiae* samples, three drops of 2 μL were placed on a CaF_2_ slide (grade 11 mm diameter by 0.5 mm polished disc, Crystran Ltd.). In each drop, 65 points were measured using a WITec Alpha300R+ with a 785nm excitation diode laser (Topotica) and a 100x/0.9 NA objective (Nikon) with 40 s of exposure and 1 accumulation using a 300 -mm/g grating.

For the samples from *C. necator, M. extorquens, Y. lipolytica* and *K. phaffi,* ~450 points were measured using 5 sec of exposure and 1 accumulation with a 300 -mm/g grating.

As a control for the instrument performance, a silica gel slide was measured with a grating of 300 - mm/g, with a 1 s time exposure and 10 accumulations. Laser power was monitored to detect possible variations. More information can be found in the Raman metadata aid (see Table S1) collected following the guidelines from García-Timermans (2018).

### Data analysis

The data analysis was conducted using R (R version 3.6.2, R Core Team 3.6.2, 2019) in RStudio version 1.2.1335 (RStudio team, 2019). Plots were produced using the package *ggplot2* and *ggpubr.* (Kassambara, n.d.; Villanueva et al., 2016).

#### Pre-processing

We manually eliminated the spectra that contained cosmic rays. The remaning spectra were preproecssed using the R packages ‘MALDIquant’ (v1.16.2)(Gibb & Strimmer, 2012) or ‘HyperSpec’ (Beleites & Sergo, 2012). To reduce the noise in the spectra, we smoothed it using the *spc.loess*() function. The 400-1800 cm^-1^ region of the spectrum (which contains the biological information in bacteria) was selected for fingerprint. The baseline was corrected for instrumental fluctuations or background noise using the Sensitive Nonlinear Iterative Peak (SNIP) algorithm (using ten iterations) and spectra were normalized using the Total Ion current (TIC). Then, the spectra were normalized using the *calibrateIntensity()* function and aligned per group with the *alignedSpectra()* function. These pre-processed data were used to calculate the single-cell phenotypic diversity and principal coordinate analysis.

#### Single-cell phenotypic diversity calculation (sc-D_2_) for single cells with Raman spectroscopy

The Hill equations were adapted in this manuscript to quantify the phenotypic diversity of single cells using pre-processed Raman spectra. Every Raman signal corresponds to a single or multiple metabolite(s), that we have called components (*n*). The relative abundance of each component was normalized, by calculating their relative abundance. Then, they were used in the Hill equation as described in the Results section.

Hill numbers are commonly used to calculate microbial diversity based on 16S rRNA gene sequencing techniques but have also been applied to flow cytometry yielding similar results (Props et al., 2016; Wanderley et al., 2019). Although there are many definitions of alpha diversity, Hill numbers are widely used. They are also known as the effective number of species, as they express in intuitive units the number of equally abundant species that are needed to give the same value of the diversity measure. Hill numbers respect other important ecological principles, such as the replication principle, that states that in a group with *N* equally diverse groups that have no species in common, the diversity of the pooled groups must be the *N* times the diversity of a single group. The general Hill equation is:

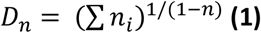

Where *q* is the sensitivity parameter, known as the order of diversity, that can be 0, 1 or 2. The diversity index of order 0 (D_0_, when q=0) corresponds to the species richness (is insensitive to the species abundance), D_1_ measures all species by their abundance, and D_2_ considers both richness and abundance.

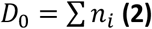

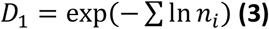

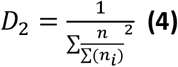

More information on the diversity measures used in microbial ecology and the advantages of Hill numbers can be found in Chao et al., 2014 and Daly et al., 2018.

#### Statistical analysis

Normality was studied using *ggdensity()* and *ggqqplot()* from the package ‘ggpubr’.

Statistics on the phenotypic diversity (sc-D_2_) of ethanol and the control group over time was done using ANOVA with the function *aov()*and post-hoc testing was done using Tukey_HSD(), both functions from the package ‘stats’.

The expression of the biomolecules in the two *S. cerevisiae* subpopulations was analysed using Wilcoxon test with the function *wilcox.test()* from the package ‘stats’.

#### Principal coordinate analysis (PCoA)

The principal coordinate analysis (PCoA) was calculated as the eigenvalues divided by the sum of the eigenvalues.

#### Sampling size

We used a dataset of 4 axenic cultures (described in table 1) and measured ~450 Raman spectra per sample, for which we calculated their single-cell phenotypic diversity (sc-D_2_). Then, we did 1000 simulations were the data were permuted, and calculated the average D_2_ when using a increasing number of spectra. The average and standard deviation of these 1000 simulations were plotted.

#### Subpopulation types

Subpopulation types were calculated by adapting the code from for flow cytometry data. The method was originally intended to separate sample clusters, while in its application for Raman spectroscopy we aim to identify and differentiate cell clusters (Props et al. 2016).

First a PCA is performed to reduce the dimensionality of the data. A reduced dataset with the principal components that explain the majority of the variance (>40%) are used to calculate the optimal number of clusters using the silhouette index, calculated with the *pam()* function from the package ‘cluster’. Once every cell is assigned to a phenotype, the median phenotype to which the (sub)population corresponds to is calculated.

#### Data availability

The analysis pipeline and the raw data can be found in https://github.com/CMET-UGent/Raman_PhenoDiv

## Results

### Phenotypic diversity quantification of Raman spectra using Hill numbers

Single-cell phenotypic changes can be captured by Raman spectroscopy, by which information is collected on the (bio)molecules present in individual cells. Once the Raman spectra are acquired, the raw data need to be pre-processed (**Fig 2-Pre-processing**). This step aims to remove noise from spectra and to be able to extract meaningful biological information. First, the spectra that contain cosmic rays need to be removed manually or automatically (Wahl et al., 2020). Then, we select the spectral region that is most relevant for microbial fingerprinting, around 500-2000 cm^-1^ (Huang et al., 2010). Once this region of the spectra is selected, the first step in the pre-processing is to correct the baseline, that can be degraded due to instrument fluctuations or background-signal influence (Liu et al., 2015; Wahl et al., 2020). Then, the spectra are normalized to avoid that the absolute intensity masks the variation of signals of interest (Beattie et al., 2009; Gautam et al., 2015). It is also possible to align and/or smooth the Raman signal, but these steps can introduce noise to the measurements and should be carefully considered.

**Fig 2:**
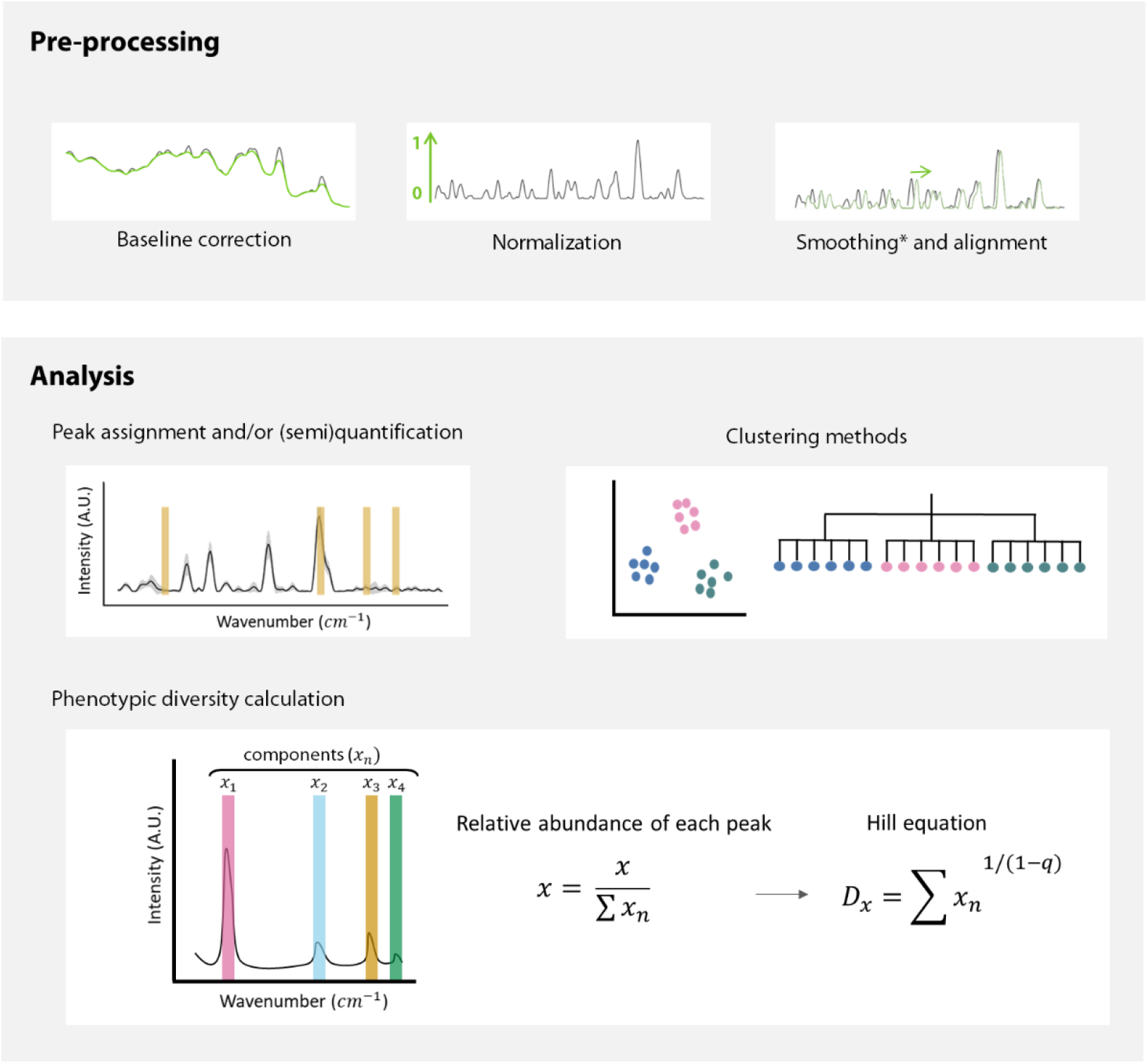
Summary of the pre-processing and analysis of the Raman spectra. First, the baseline is corrected, and the spectra are normalized. Spectra can be smoothed and aligned; however, smoothing can erase potentially relevant information, and should be carefully considered. Similarly, alignment can produce faulty spectra by displacing the signal, and thus need to be used reasonably. Once the spectra are pre-processed, it is possible to (1) extract (semi)quantitative information (2) cluster cells or create phenotypic trees or (3) calculate the single-cell phenotypic diversity. For the latter, Raman peaks that correspond to one or several metabolites are considered as components. The intensity of these components (*x*) is used to quantify phenotypic diversity. The order of diversity (*q*) can be 0, 1 or 2, meaning respectively that richness, abundance or both parameters are considered in the metric. This equation considers richness and estimated abundance of metabolites in a single cell.

After the spectra have been pre-processed, different information can be extracted (**Fig 2-Analysis**). For example, peaks of interest can be selected for semi-quantitative analysis or quantitative analysis using a calibration curve (Butler et al., 2016). Also, the whole spectra can be used to classify cells using several clustering methods, such as principal component analysis, principal coordinate analysis, non-metric multidimensional scaling or T-distributed stochastic neighbour embedding. This information can also be used to construct phenotypic trees (Garcia-Timermans 2018). Here we used the preprocessed spectra to quantify the single-cell phenotypic diversity using Hill numbers. Every Raman peak corresponds to a different metabolite or a combination of metabolites, called components (*x*) (**Fig 2**). To calculate the relative abundance of each peak, the intensity of the signal of each component was normalized by the sum of all intensities, and this information was then used in the Hill equations.

The order of diversity (*q*) can be 0, 1 or 2, meaning that richness, abundance or both richness and abundance are taken into account in the metric. sc-D_0_ contains information about the number of components *(n_i_)* in the Raman spectra, and is calculated as shown in equation 2.sc-D_1_ informs about the abundance of each component and is described in equation 3. In this paper, we mostly focus on single-cell D_2_ (sc-D_2_) (*q*=2) as it takes both richness and abundance of the Raman components into account.

### Sample size dependence of phenotypic diversity (sc-D_2_) measurements

To understand the distribution of single-cell phenotypic diversity in a population, we did ~450 measurements in 4 axenic cultures of *C. necator, M. extorquens, Y. lipolytica* and *K. phaffi.* We calculated the average diversity estimation for an increasing number of spectra and bootstrapped 1000 times. The average of the total number of measurements is plotted in grey, and the 5% of this average is represented with a dotted grey line.

We looked at how many mreasurements were needed to calculate the population average (grey line) and how many are needed to have an accurate estimation (95%, dashed lines). For the estimation of sc-D_0_, few measurements (~10-50) are were needed to obtain the population average. The sc-D_1_ calculation grants a greater weight to high-intensity wavenumber and/or peaks of these components, and required ~100 measurements. Although *M. extorquens* reaches it after ~20 measurements. The sc-D_2_ estimation takes both the number of components and their abundance into account and needed between ~50 (*C. necator)* to ~180 (*Y. lipolytica)* measurements to estimate the population average.

### Case studies: phenotypic diversity quantification in stress-induced phenotypes

When stress is applied in a microorganism, a set of genes and proteins are expressed, changing the metabolic phenotype of the cell. This metabolic change can be captured by Raman spectroscopy, that collects information on the (bio)molecules present in individual cells. To compare stressed and nonstressed cells, we quantified their phenotypic diversity using our proposed methodology, as shown in **Fig 1**. First, we compared two *E. coli* cultures growing in different conditions: with ethanol (stressed) or non-treated (control). Then, we compared two subpopulations of the same *S. cerevisiae* culture, separated based on their expression of the GFP stress reporter in nutrient-limiting conditions.

#### Tracking *E. coli* population diversification dynamics following exposure to ethanol stress

We used a dataset from Teng 2016, consisting of spectra of *Escherichia coli* sampled at different time points (5, 10, 20, 30 and 60 min, 3 h and 5 h) after being cultured in standard conditions or with ethanol. There were three biological replicates of the cell culture and 20 cells were measured per replicate.

The stress-induced metabolic diversity of single cells was quantified using the sc-D_2_ Hill equation and the average diversity for each population (stress and non-stressed) was plotted (**Fig 4A**). After testing for normality, a two-way ANOVA test showed a significant difference between treatments and treatments over time (p < 0.0001). A post-hoc Tukey test showed that the ethanol and control groups were significantly different at time point 60 min and 180 min (p < 0.0001). Then we used PCoA, a common clustering method to visualize the dissimilarities in the fingerprints. The Raman fingerprint of the stressed and control cells is similar at the beginning and then shift over time (**Fig 4B**). We used a clustering algorithm to define exactly when this shift takes place: after 20 min for the ethanol-treated population and 180 min for the control population (**Fig 4C**).

**Fig 3:**
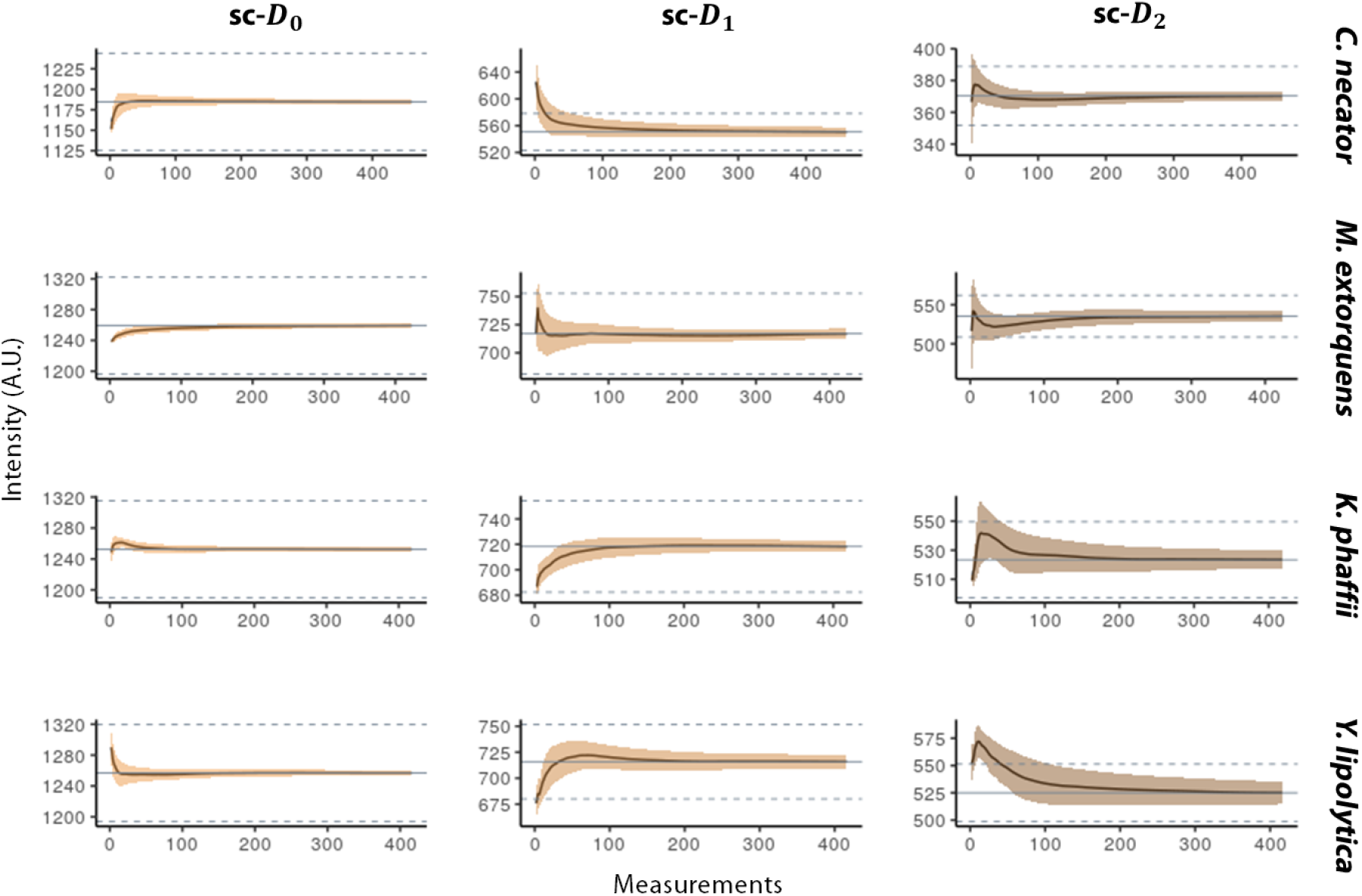
Effect of sampling size on the single-cell phenotypic diversity average. We calculated the average single-cell phenotypic diversity using the Hill equations (single-cell D_0_, D_1_ and D_2_) for an increasing number of measurements and repeated the calculation picking spectra randomly 1000 times. We used the Raman spectra of four pure cultures and ~450 measurements on each. The smear represents the standard deviation. The grey line represents the average sc-D value of the total population, and the dashed lines a 5% deviation from the mean.

**Fig 4:**
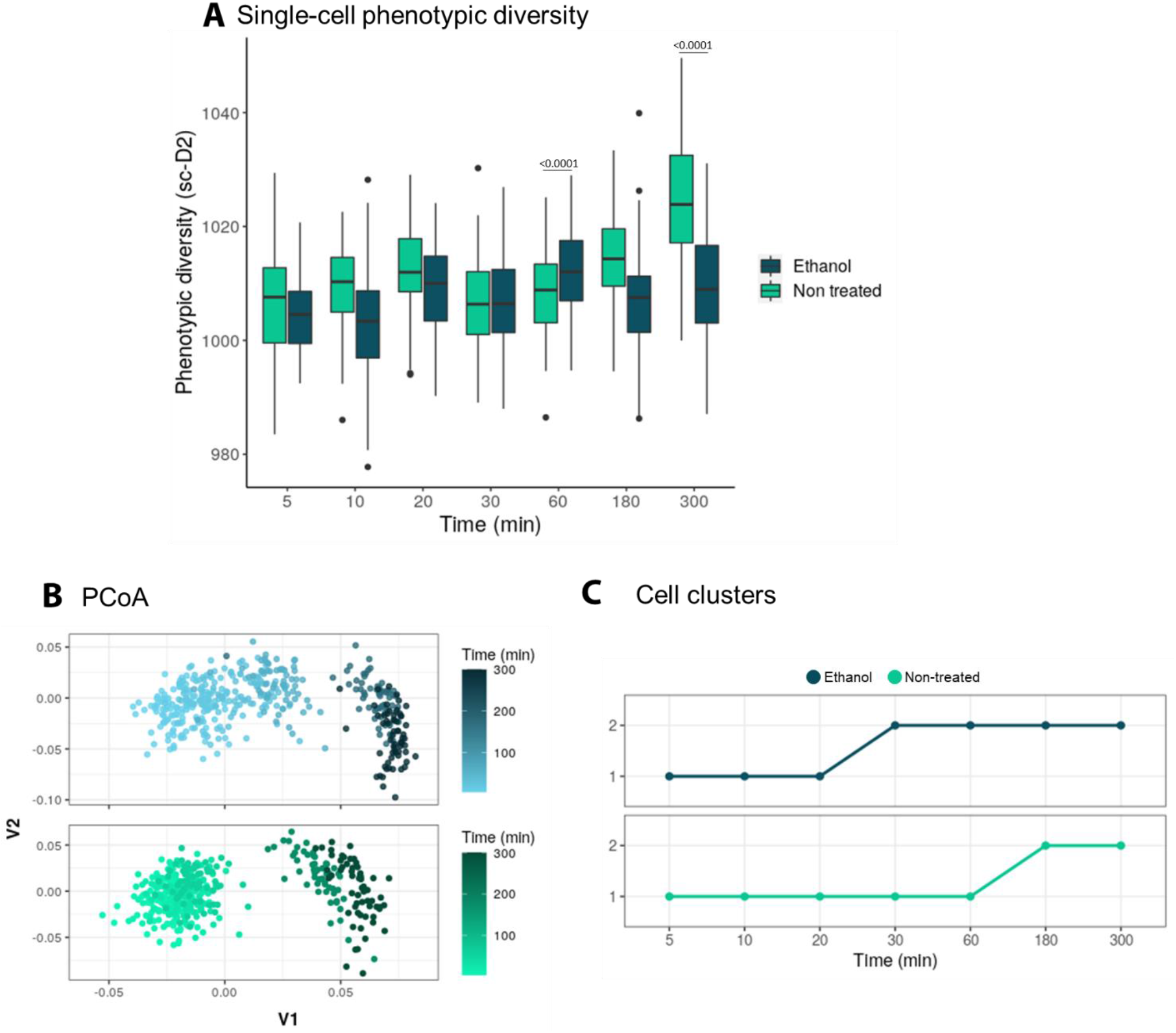
**A)** Single-cell phenotypic diversity (sc-D_2_) of the stressed (ethanol treated) and non-stressed (non-treated) *E. coli* populations. Treatments and treatments over time are significantly different (two-way ANOVA, p < 0.0001). A post-hoc Tuckey test showed that the ethanol and control groups are significantly different on timepoint 60 min and 180 min (p < 0.0001). **B)** Raman fingerprint of the stressed (ethanol treated) and non-stressed (non-treated) *E. coli* populations, plotted using principal component analysis (PCoA). The time progression is represented with a darker colour. Every point represents a single cell. **C)** The clustering algorithm shows the phenotypic shift happens after 20 min for the ethanol-treated population and after 180 min for the control. Two phenotypes were found. Every point represents the average “phenotypic type” of the population. N=60

#### Discriminating *S. cerevisiae* subpopulations following exposure to nutrient limitation

A *S. cerevisiae* population was cultured in nutrient-limiting conditions. Based on GFP expression as an indicator of stress activation, we separated two subpopulations (one that activated the stress reporter, and one that did not) using FACS. Then, we analyzed 65 cells in each subpopulation using Raman spectroscopy.

First, we calculated the single-cell phenotypic diversity (sc-D_2_) of the subpopulations with high (+) or low(-) stress reporter expression. To prove that sc-D_2_ calculations are quantitative, we also created an *in silico* group by mixing the two subpopulations (**Fig 5A**). The *in silico* mix group was expected to have an average sc-D_2_. Then, we checked the dissimilarity of the fingerprints using PCoA (**Fig 5B**). Two clusters are differentiated depending on the reporter expression.

**Fig 5:**
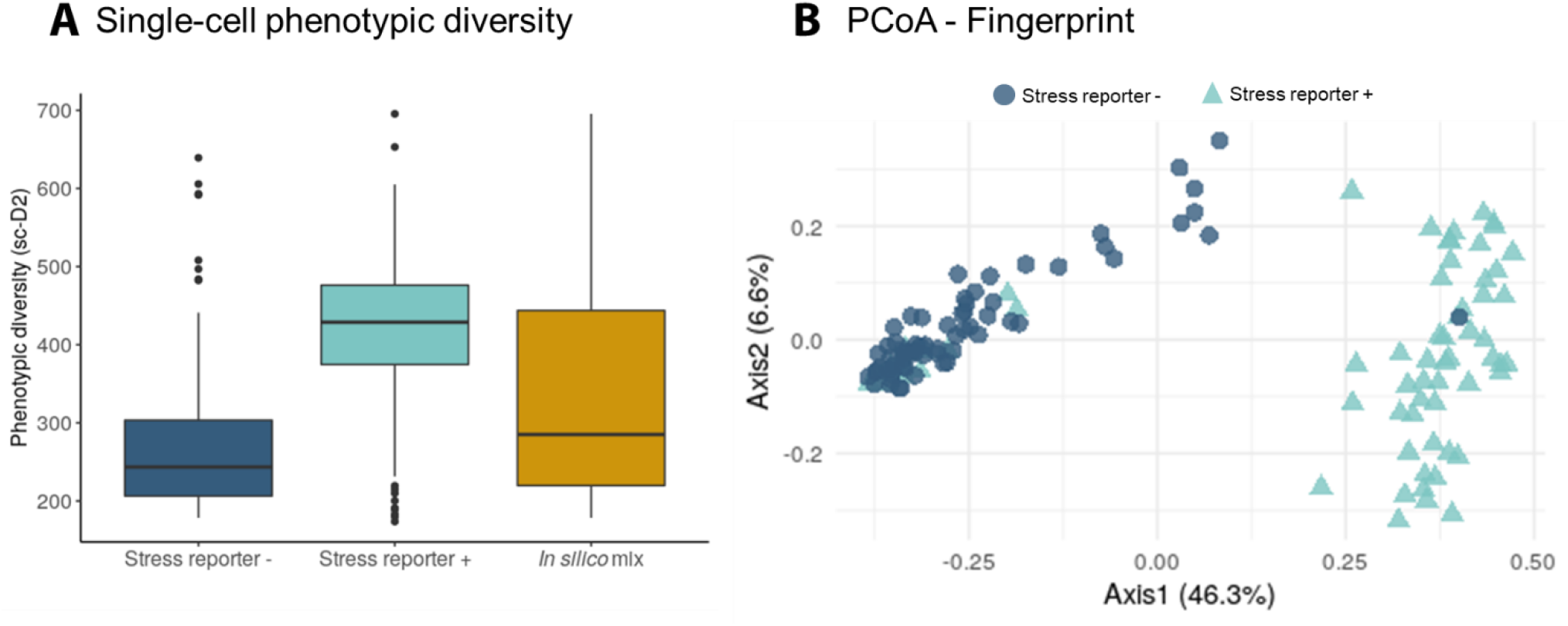
**A)** Single-cell phenotypic diversity of a *S. cerevisiae* subpopulations with high or low stress reporter expression and an *in silico* mix of both groups. The *in silico* mix is a random selection of cells coming from the stressed and non-stressed population **B)** Visualization of the stress-induced phenotypic change of *Saccharomyces cerevisiae* subpopulations with high or low stress reporter expression using principal coordinates analysis (PCoA). Every dot is a single cell. The size of the dot corresponds to the single-cell phenotypic diversity (sc-D_2_). N= 65.

The information of the Raman spectra from each group was used to understand the effect of the stress reporter activation on the metabolic response of *S. cerevisiae.* The intensity of the Raman peaks that correlate well to the content in proteins, lipids, nucleic acids and unsaturated lipids (Teng *et al.,* 2016) was compared in the subpopulations with a high or low stress-reporter expression (**Fig 6**). We found that both groups have a significantly different metabolism: the subpopulation with a high (+) expression of the stress reporter had more unsaturated lipids and proteins, but contained less lipids and nucleic acids (Wilcoxon rank-sum test, p < 0.0001).

**Fig 6:**
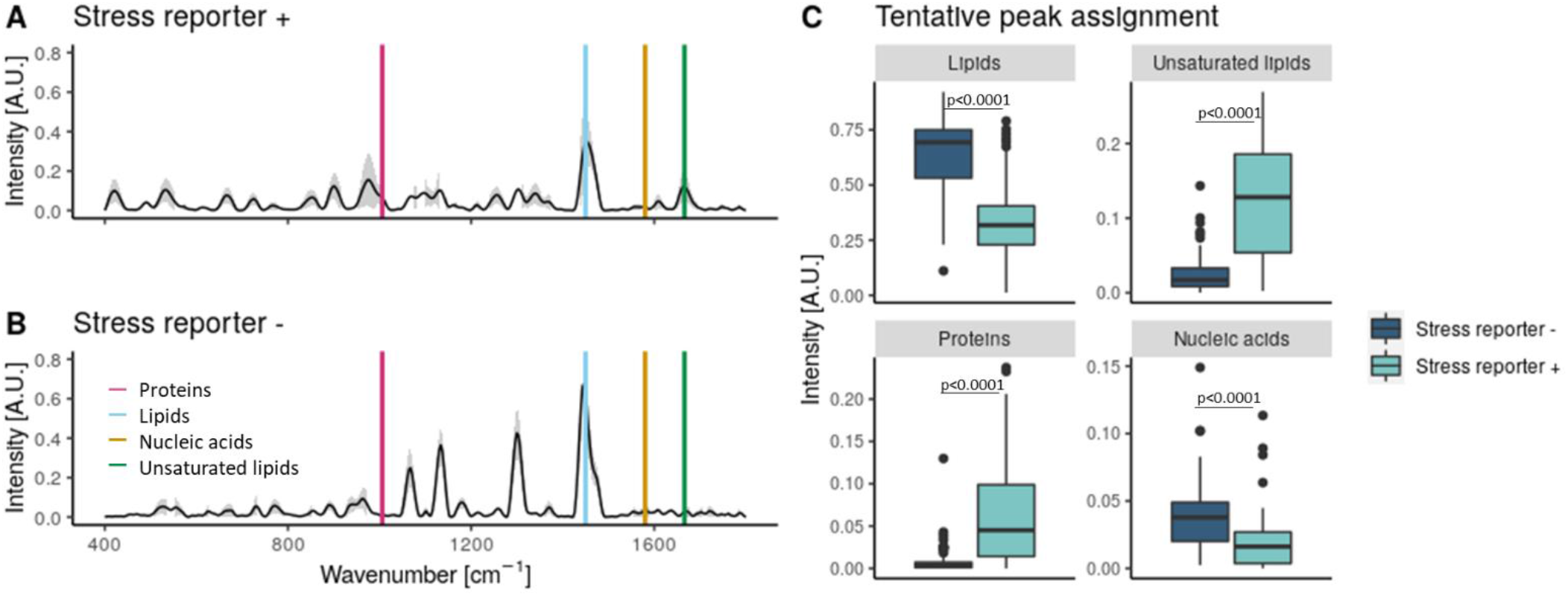
Raman spectra of *S. cerevisiae* subpopulations with high (+) or low (-) expression of the stress reporter **(A,B)**. The average of the spectra is plotted with a black line and the standard deviation in grey. The putative peaks corresponding to proteins, lipids, nucleic acids and unsaturated lipids according to Teng and colleagues (2016) are plotted over the spectra. **C)** The intensity of the metabolic peaks highlighted in plot A and B for the subpopulations with high or low expression of the stress reporter. The *p* values for the Wilcoxon test for every metabolite is shown. N=65

## Discussion

Raman spectroscopy can quantify stress-driven metabolic heterogeneity at the single cell level to detect how and when bacteria diversify their metabolism. This tool is relatively fast and non-destructive, and can provide (semi)quantitative information about the composition of cells. Once the spectra are measured, they need to be pre-processed to remove as much noise as possible. First, the samples with cosmic rays can be removed manually, or the cosmic rays can be subtracted automatically. Then the baseline is corrected, and spectra are normalized (**Fig 2**), although there is some discussion as to whether these calculations should be performed in a single step (Liu et al., 2015; Wahl et al., 2020). There are other possible data transformations, such as the aligning the spectra, to avoid the small instrumental variations that can show up (García-Timermans et al., 2018). However, this step might introduce noise (i.e. by misplacing Raman signals) and should be carefully considered. Smoothing can be also used, but this step can erase small points in the spectra, removing relevant information. In our case, we noticed the spectra were noisy, and decided to smooth the spectra. We also aligned the samples per group, although this had very little effect in the dataset.

Once the spectra have been pre-processed, they can be used to investigate the phenotypic heterogeneity among or within cells. Although Raman spectroscopy has been previously used to detect stress-driven phenotypes (Tanniche et al., 2020), we argue that there is a need for single-cell quantitative measurements for phenotypic diversity and propose the use of Hill numbers. We chose Hill numbers for our calculations because they are widely used in microbial ecology. As previously stated, they are easy to understand because they represent the effective number of species - the number of equally abundant species needed to give the same value of diversity measure – and respect important ecological principles such as monotonicity in the number of species and the replication principle (Daly et al., 2018).

To estimate phenotypic diversity using Hill numbers, we considered that each Raman signal corresponds to a component (a single or multiple molecules), and that the intensity of these components is correlated with their quantity (Tang et al., 2013; Wu et al., 2011). After normalizing the components, they were used in the Hill equations (**Fig 2**). Although we chose to use the whole spectrum for this calculation, it is possible to select only the peaks. However, this could influence the resolution: algorithms for peak detection typically divide the spectrum according to a certain window size and look for the local maximum (Gibb & Strimmer, 2012). Using this algorithm would not take into account the width of components, which is a characteristic of the molecules. Also, some components with a close signal would be ignored, and the choice of window size would affect the final result. How the Raman spectra are preprocessed will have an impact on the results. The region used for fingerprinting needs to be considered so that all the relevant biomolecules to address the hypothesis are reported. Both the baseline correction and normalization will have an impact on the intensity reported for the different components. Smoothing functions assume spectra are noise, and erase certain signal. Finally, aligning spectra when unnecessary can misplace the signals. Using the same preprossing steps when comparing samples is crucial, as well as detailing the preprocessing steps and providing the raw data.

To explore the importance of the sample size in these estimations, we used a large dataset consisting of ~450 Raman spectra from 2 axenic bacterial cultures (*C. necator* and *M. extorquens)* and 2 axenic yeast cultures (*Y. lipolytica* and *K. phaffi).* Then, the effect of the sampling size on the average singlecell phenotypic diversity and its standard deviation was calculated. Our results show that this is highly population-dependent: for example, while *C. necator* only needed 15 spectra to approach the expected *sc-D_2_* average, *Y. lipolytica* needed more than 150 measurements (**Fig 3**). This could be due to a different degree of phenotypic diversity in the populations. Sample size should be explored for every experiment, to make sure that the estimations are representative.

After developing the methodology to quantify single-cell phenotypic diversity, we applied it to two case studies to demonstrate its use. We focused on sc-D_2_, as it considers how many components are being expressed per cell, and their abundance. In the first case study, we compared an ethanol-treated and a control *E. coli* population. We found that when *E. coli* is grown in standard conditions, there is a phenotypic shift after 60 min. This shift happens earlier in stressed cells (20 min) (**Fig 4C**). The shift in the fingerprint in the control group could be due to the entering in the log phase. Our group previously showed how *E. coli* start their log phase after ~1h of cultivation in rich medium, and how at different growth stages bacteria change their phenotype (García-Timermans et al., 2019). Although both the ethanol-treated and the control populations end up having a similar phenotype after 60 min, the stressed population has a lower metabolic diversity (**Fig 4A**), a lower nucleic acid content and a higher protein and lipid content. Clustering algorithms are useful to automatically identify phenotypes and quickly asses when the phenotype of a population has changed in a reproducible way. While here we use PCA, other metrics can be used, such as non-metric multidimensional scaling (NMDS), t-distributed Stochastic Neighbor Embedding (t-SNE) and other clustering methods. The choice of the clustering method should be based on the hypothesis, and how important it is to conserve the distances between the cells and the relative size of the cluster.

In the second case study, we analyzed the response of two *S. cerevisiae* subpopulations. When in nutrient-limiting conditions, *S. cerevisiae* resorts to a bet-hedging strategy where some yeasts will enter a quiescent state, while others will activate a stress-induced response (Gray et al., 2004). The strain used in this experiment produces GFP upon activation of nutritional stress, so when the *S. cerevisiae* culture diversified into two populations -with either high or low expression of the stress reporter-these were separated using FACS and analyzed with Raman spectroscopy. Because the Raman spectroscope used has a 785 nm laser, we do not expect the fluorescent signal (excited at 510 nm) to be picked up with this instrument. Single-cell phenotypic diversity (sc-D_2_) in the stressed subpopulation is higher than the non-stressed (**Fig 5A**). As expected, the *in silico* mix shows a diversity that is close to the average of both subpopulations. We then checked that the subpopulations with high and low stress reporter expression had a different fingerprint using PCoA, a tool widely used for Raman spectra in microbial ecology. This confirmed that the fingerprint of both subpopulations is visibly different (**Fig 5B**). Using the metabolic information contained in the Raman spectra, we found a higher nucleic acid content in the non-stressed subpopulation (in line with the findings of Teng 2016 in stressed *E. coli* cells). This could be explained by the higher ribosome content in non-stressed cells. We also found that the stress response triggered by the activation of the chimeric promoter results in a raise of protein and unsaturated lipids production (**Fig 6**), similar to the results found in stressed *E. coli* cells. However, it could be that the protein responsible for this difference is (at least partially) the GFP protein itself. The choice of this promoter based on a fusion of *glc3* and *hsp26* as a single proxy to define a metabolically stressed population is cross validated by these findings, that show two clearly metabolically distinct subpopulations.

Finally, we explored whether the number of cells measured in both case studies was enough to capture the diversity of the cultures. In *S. cerevisiae,* 65 cells were enough to estimate single-cell diversity, and most biomolecules **(Fig S2, Fig S3)**. However, to properly estimate the protein content in the non-stressed subpopulation more cells would have been needed. In the *E. coli* population, we tested the sample size in the ethanol-treated population at timepoint 5 min and 300 min. Very few cells are needed to have a representative single-cell diversity estimation: the sc-D_0_ is the same for all cells (**Fig S4**). This metric looks at the number of components present in each cell, which in this case seem to be the same for all individuals. It could be that these cells express the same molecules, but different amounts, and/or an artefact of the pre-processing carried out by Teng *et al,* that could have erased some of the smaller peaks. This highlights the importance of making the raw data available, following the trends of other disciplines such as new generation sequencing (NGS) or flow cytometry.

Inferring metabolic expression from Raman spectra in microbial cells is not without challenges. For instance, many databases propose different peaks to identify the same biomolecules. In this manuscript, we have chosen those presented in Teng et al. 2016 to be able to compare the results they found in *E. coli* and we found in *S. cerevisiae.* Some molecules are not Raman active, and thus will not be reflected in the spectra. Conversely, some Raman active molecules can be overrepresented in the analysis. Also, there can be Raman peaks that correspond to several compounds. These limitations should be considered when using Raman spectroscopy for microbial ecology. A better assignment of the Raman signals will also contribute to an improved understanding of the metabolic changes driving single-cell phenotypic heterogeneity.

Raman spectroscopy is a promising single-cell technology, able to quantify phenotypic diversity in individual cells, identify changes in phenotypes and estimate metabolic information (semi)quantitatively. Single-cell tools represent the next challenge of microbial ecologists: they can go beyond community measurements, based mostly on single marker-gene expression or low-dimensional physiological data, and shed light on how heterogeneity shapes communities.

## Conclusions

- Raman spectroscopy can be used to quantify single-cell stress-driven phenotypic diversity in microbial communities.
- Each Raman spectral point corresponds to a different metabolite (or to multiple metabolites), that are expressed with a certain abundance (intensity). Using this information in the Hill diversity framework, we can estimate the phenotypic diversity in single cells. We show that these methods work to study changes at the population and subpopulation level in both prokaryotes and eukaryotes.
- The Raman spectra contain information about the biomolecules present in a cell, and can be used to study the metabolic shift in stressed cells.
- We propose an automatic classification of phenotypes using clustering methods. This is a useful tool to track changes in singe-cell physiology.

## Declarations

### Data availability

The raw data and code to reproduce the analysis shown in this manuscript can be found in the repository https://github.com/CMET-UGent/Raman_PhenoDiv

The dataset from Teng et al. 2016 was used to validate alpha and beta-diversity calculations, as well as the ‘subpopulation type’ definition.

### Author contributions

CGT wrote the paper with contributions from RP, BZ, FD and NB. BZ and FD cultivated, harvested and sorted the *S. cerevisiae* cells. MS cultivated and harvested *C. necator, M. extorquens, Y. lipolytica* and *K. phaffi.* CGT collected the Raman data. CGT performed the data analysis with the help of RP. CGT, RP, BZ, FD and NB designed the study. All authors read and approved the final version of the manuscript.

## Acknowledgements

The authors thank the funding that made this research possible. CGT is funded by the Flemish Fund for Scientific Research (FWO G020119N) and by the Geconcerteerde Onderzoeksacties (GOA) research grant from Ghent University (BOF15/GOA/006). RP is supported by the Flemish Fund for Scientific Research (FWO). BZ is supported by a post-doctoral grant through an Era-Cobiotech project (“ComRaDes” Computation for Rational Design: From Lab to Production with Success). MS is supported by the Catalisti cluster SBO project CO2PERATE (“All renewable CCU based on formic acid integrated in an industrial microgrid”), with the financial support of VLAIO, Belgium (Flemish Agency for Innovation and Entrepreneurship). This project has received funding from the European Union’s Horizon 2020 research and innovation programme under grant 722361.

